# Aβ_42_ Facilitates the Activation of Discoidin Domain Receptor 2 and its Nuclear Enrichment in Alzheimer’s Disease Model

**DOI:** 10.64898/2025.12.16.694181

**Authors:** Russa Das, Sampa Biswas, Debashis Mukhopadhyay

## Abstract

Discoidin domain receptor 2 (DDR2), a receptor tyrosine kinase (RTK) family member, plays a pivotal role in collagen-mediated signaling, influencing several key cellular processes. In Alzheimer’s disease (AD), DDR2 expression and activity are significantly upregulated. While collagens, the endogenous ligands of DDR2, either remain unchanged or are downregulated in AD-like conditions, Amyloid-beta 42 (Aβ_42_) has been found to activate DDR2 in the absence of collagen non-canonically. Additionally, DDR2 naturally translocates to the nucleus in the neuroblastoma cell line SH-SY5Y, and the pool of DDR2 increases further in the nucleus in the presence of Aβ_42_, suggesting its novel pathological role in AD. The E113K mutant of DDR2, where the collagen-binding site has been compromised, fails to get activated by collagen, whereas Aβ_42_ could still activate it. Furthermore, the DDR2 inhibitor WRG-28 effectively inhibits Aβ_42_ induced phosphorylation of the protein.

We have also modeled *in silico* the ectodomain of DDR2, extending through the transmembrane helix, and subsequently docked Aβ_42_ pentamer to elucidate the mechanistic basis of kinase activation by amyloid Aβ. Additionally, we analyzed WRG-28 binding by constructing the ternary complex of DDR2, Aβ_42_, and WRG-28. Our structural analyses offer key insights into the molecular mechanisms governing DDR2 activation by Aβ_42_ and its inhibition by WRG-28 in the AD cell model, providing a foundation for targeted therapeutic strategies.

**Graphical Abstract:** 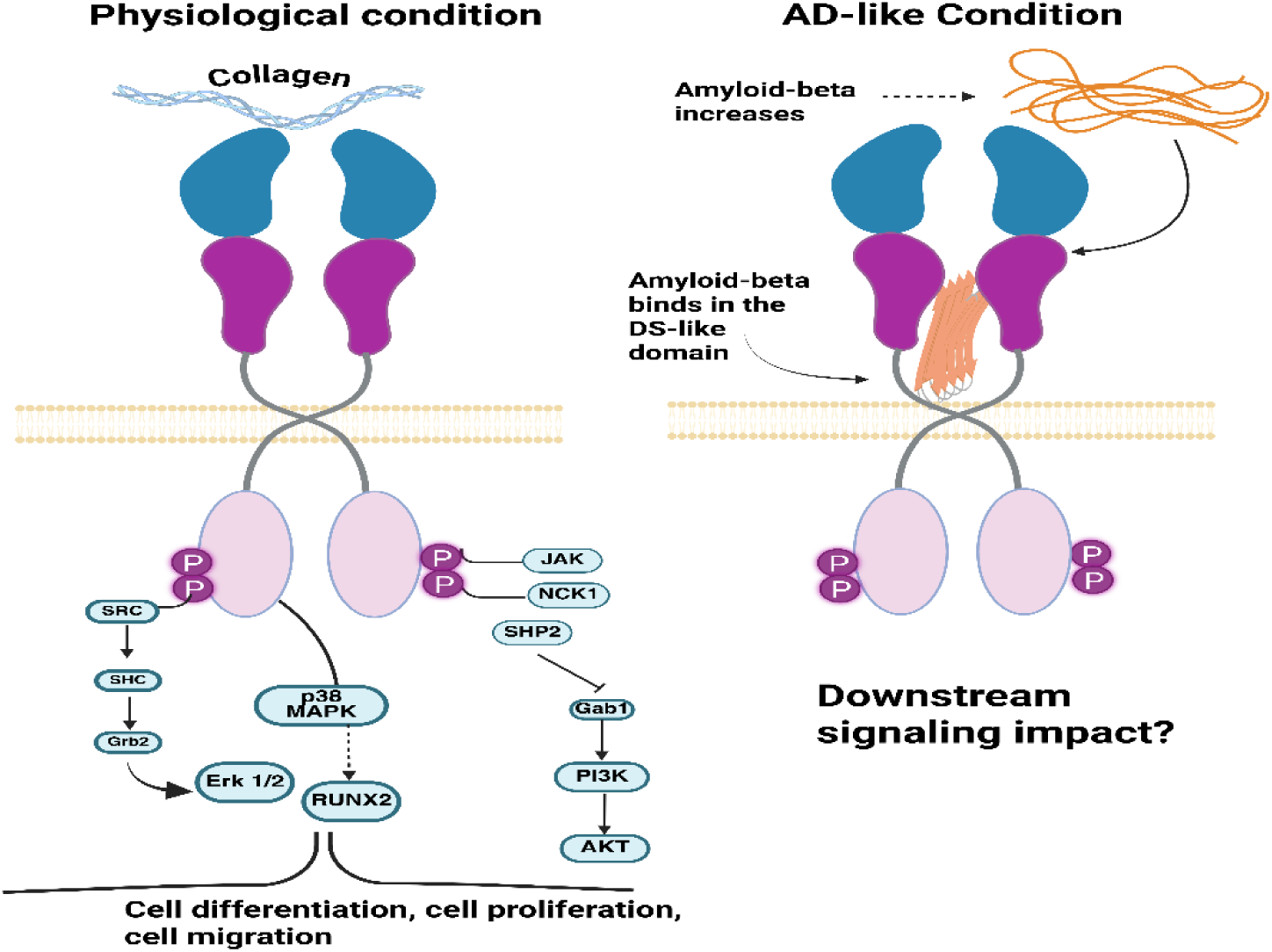

## Introduction

Discoidin Domain Receptors (DDR) belong to a unique Receptor Tyrosine Kinase (RTK) family that binds to its ligand collagen and mediates downstream signaling ^1,2^. DDR consists of an extracellular region that includes the N-terminal Discoidin (DS) domain, a DS-like domain, the extracellular juxtamembrane (JM) domain, and a large transmembrane (TM) region, followed by the cytosolic catalytic domain with a short C-terminal tail ^3^. The two members DDR1 and DDR2 exhibit substantial sequence similarity, with 59% identity in DS domains and 51% in DS-like domains^4^. While DDR1 is mainly activated by fibrillar collagens I and V and non-fibrillar collagen IV, DDR2 is exclusively activated by fibrillar collagens (types I-III and V) ^5^. DDRs are widely expressed in diverse tissues during developmental stages as well as in adults. DDR1 mRNA is detected in both murine and human subjects, with elevated levels in the brain, lung, kidney, spleen, and placenta. DDR2 mRNA is highly expressed in skeletal, cardiac, kidney, and lung tissues. Notably, both are present in the developing nervous system, indicating their roles in neural development. DDR1 is predominantly expressed in epithelial cells, while DDR2 is mainly found in connective tissue cells originating from the embryonic mesoderm^4^. A unique feature of DDRs is their slow activation mechanism. In contrast to the rapid interaction of soluble growth factors with receptor tyrosine kinases, collagen binding to DDRs triggers slow autophosphorylation of the receptor tyrosine, taking approximately 60 to 90 minutes to achieve full activation and lasting for up to 18 hours^6–8^. Tyrosine residues 736,740,741 located in the DDR2 kinase domain upon phosphorylation regulate the kinase catalytic activity^6^. Additionally, *in-vitro* kinase assays with DDR2 demonstrated that Tyr740 and Tyr741 are phosphorylated only when DDR2 achieves its maximum kinase activity^9^. Similarly, Collagen binding to DDR1 triggers slow autophosphorylation of multiple tyrosine residues within the activation loop, including Tyr792, Tyr796, and Tyr797^10^. Phosphoproteomic analysis revealed that insulin enhances collagen I signaling by increasing DDR2 phosphorylation which may have significant implications for DDR2’s role in health and disease^11^. Under normal physiological conditions, DDRs perform a wide range of cellular functions including cell migration, proliferation, and extracellular matrix remodeling ^6^, whereas loss-of-function mutation of DDR2 in humans and mice caused serious defects in skeletal growth and development^12^. Dysregulation of DDR signaling has been associated with various diseases including cancer, atherosclerosis, and fibrotic disorders^13^.

DDR’s involvement in neurodegeneration, however, remains poorly understood. Especially for AD, the formation of insoluble amyloid β-peptide (Aβ) has been a major histopathological hallmark ^14^. An RTK activity assay in the presence of Aβ_42_ oligomers showed that out of 58 known human RTKs, 18 RTKs were differentially regulated in AD, both DDR2 and DDR1 being the significant ones among them^15^. Both DDR1 and DDR2 levels were upregulated in *post-mortem* Alzheimer’s and Parkinson’s brains, and lentivirus-mediated knockdown of these two receptors reduced the levels of α-synuclein, tau, and β-amyloid proteins^16^. In a mouse model challenged with α-synuclein, partial or complete deletion or inhibition of DDR1 promoted autophagy, suppressed inflammation, and decreased the accumulation of neurotoxic proteins^13^. Several pieces of evidence also suggest that the Aβ can bind several molecules and receptors on the neuronal surface and mediate a downstream signaling cascade contributing to AD pathology^17^. These receptors include the prion protein (PrPC), N-methyl-D-aspartate receptor (NMDAR), the receptor for advanced glycation end products (RAGE), and the p75 neurotrophins receptor (p75NTR), besides the RTKs. Upon binding, Aβ induces synaptic dysfunction, disrupts calcium homeostasis, and promotes neuroinflammation and oxidative stress. This interaction between Aβ and cell surface receptors ultimately leads to neuronal damage, loss of cognitive function - the hallmark of neurodegeneration observed in Alzheimer’s disease^18,19,20,21^. All these boil down to the potential role of DDR2 in Alzheimer’s disease, even though it has only been minimally explored.

With increasing evidence supporting the involvement of DDR2 receptors in Alzheimer’s disease, we sought to investigate how DDR2 is regulated in our AD cell model in terms of its activation and subcellular localization. DDR2 is typically found in the plasma membrane as a dimer (Fig 1A). However, recent evidence has indicated the presence of phosphorylated DDR2 (pDDR2) in nuclear-enriched fractions of Adipose-derived (AD), Bone marrow-derived (BM), and MScs derived from breast cancer metastatic cells (Met-MSCs)^22^. In this present study, human neuroblastoma cells (SH-SY5Y) were made to mimic an AD-like scenario and were used to investigate the role of DDR2. The protein expression database (https://www.proteinatlas.org/) indicates that SH-SY5Y cells have inherently low expression levels of several collagen types (Col I, II, III, V, and X), which are the primary canonical ligands of DDR2. Additionally, exposure to 5 µM Amyloid-β (Aβ_42_) for 24 hours has been reported to downregulate collagen expression in these cells^23^ .Therefore, this study aims to investigate the regulation, activation status, and subcellular localization of DDR2 in SH-SY5Y based AD-like cell model following Aβ_42_ exposure. This will provide new insights into the non-canonical activation mechanism of DDR2 in neurons and its potential contribution to AD pathology. Uncovering these mechanisms could establish DDR2 as a novel therapeutic target or biomarker in neurodegenerative diseases like Alzheimer’s disease.

**Figure 1:**
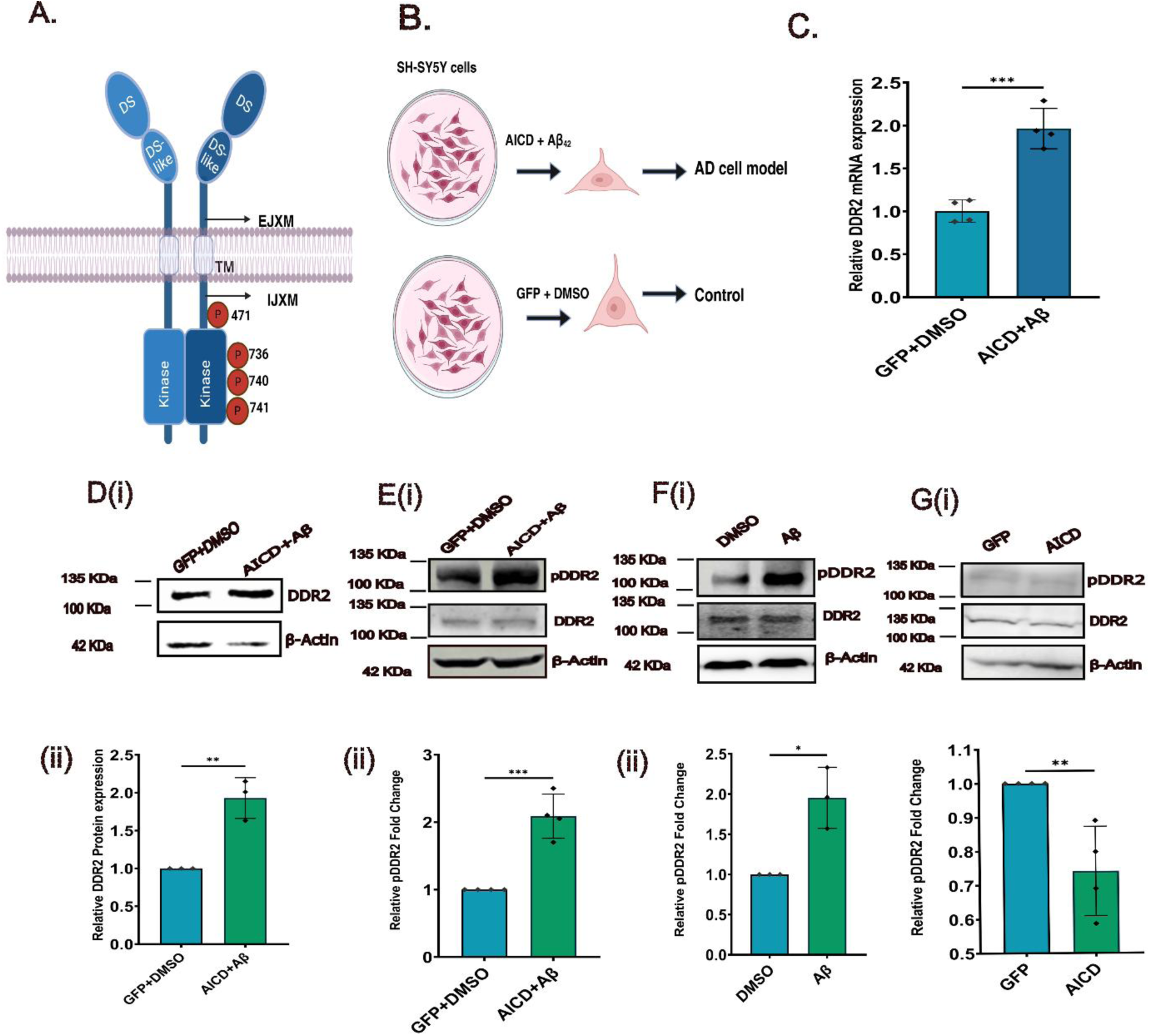
(A) Cartoon structure of DDR2 Dimer represents the DS-domain, DS-like domain, extracellular juxtamembrane domain, intracellular juxtamembrane domain, and kinase domain. All the phosphorylation sites have been marked in the kinase domain (red circles) (B) Representative of the AD-like cell model. (C) The relative fold change in the DDR2 transcript level was analyzed in the AD cell model (n=4) and normalized with GFP and DMSO. (Di, ii) DDR2 protein expression level significantly increases in the AD cell model compared to the control (n=4). (Ei, ii) DDR2 activity also significantly increases in the AD cell model (n=4). (Fi, ii) Relative fold change in the DDR2 activity in Aβ treated cells compared to DMSO (n=3). (Gi, ii) There is no significant change in the DDR2 activity in AICD-only treated cells compared to GFP as control. The error bars were calculated as mean (fold change) ± SD values. The *p*-value of ≥ 0.05 was considered to be statistically insignificant data and was denoted as “ns”. Statistical significance was represented as follows *p value < 0.05; **p value < 0.01; and ***p value < 0.001

## Results

### Expression and Activity levels of DDR2 are altered in AD-like pathological condition

We analyzed DDR2 expression at both the transcript and protein levels in the *in vitro* AD cell model. The results showed significant upregulation of DDR2 mRNA (0.96 ± 0.13) and protein expression level (0.93± 0.15) (Fig 1C, 1D). Additionally, we assessed DDR2 activation in the AD cell model and its control. DDR2 activity was significantly elevated in the AD cell model (1.128 **±** 0.27) and Aβ_42_-treated cells (1.65 **±** 0.14) compared to their respective controls, whereas no significant change was observed in AICD-GFP-transfected cells (Fig.1E,1F,1G).

To gain more insight into the enhanced activity of DDR2 in the AD cell model, we also checked the expression levels of collagens, which are the canonical ligands of DDR2. The transcript level expression of collagens either did not show any significant changes, or it was decreased in AD-treated cells as compared to the control (Supplementary Figure 1). We also investigated the individual effects of each of the pathological parameters, like Aβ_42_ and AICD, separately. In the presence of AICD-GFP, the DDR2 activity level showed no significant difference, whereas in the presence of Aβ_42,_ the DDR2 activity level significantly increased. This implied that even in the absence of collagen, Aβ_42_ interacted with DDR2 and activated it in the AD context.

### Aβ_42_ colocalizes with DDR2 in absence of collagen in AD-like conditions

Because even in the absence of collagen DDR2 was getting significantly upregulated in AD, we asked whether Aβ_42_ could interact with DDR2. To address this question, we treated the SH-SY5Y cells with Aβ_42_ for different time lengths. After 12 hours of treatment, double immunofluorescence studies showed that in Aβ_42_ treated cells where DDR2 (green) was present, Aβ42 (red) was also present. The close spatial relationship between Aβ_42_ and DDR2 in these cells was further highlighted when the two filters were overlaid (appearing yellow) suggesting a strong colocalization between Aβ_42_ and DDR2 (Fig:2A) with a Pearson correlation coefficient value of 0.657 ± 0.029. Incidentally, when looking for the interaction between these two, we observed that DDR2 also localizes inside the nucleus. This intrigued us to investigate the sub-cellular localization of DDR2 under normal physiological conditions.

### DDR2 undergoes nuclear translocation in cells of neuronal origin

We performed confocal microscopy on untreated SH-SY5Y cells using a monoclonal antibody targeting the C-terminal region of DDR2 (amino acids 422–639) to assess its subcellular localization. Immunofluorescence analysis revealed prominent nuclear localization of DDR2, which was further validated by subcellular fractionation and immunoblotting (Fig. 2B, 2C). To assess whether this nuclear localization had been unique to neuronal cells or not, we repeated the same experiment in non-neuronal HeLa cells, and no nuclear localization of DDR2 was observed (Supplementary Figure 2). To investigate Aβ_42_ whether could influence DDR2 distribution, we compared nuclear DDR2 levels in Aβ-treated cells versus DMSO-treated cells (relevant control). Aβ exposure led to a marked increase in nuclear DDR2 (Fig. 2D, 2E). Moreover, upon detailed inspection of the immunofluorescence images in Figure 2D enhanced perinuclear staining of DDR2 was observed in Aβ-treated cells indicating a potential intermediate step in its translocation or distribution towards the nucleus. This observation suggests that Aβ_42_ may not only drive nuclear translocation but also modulate trafficking or retention of DDR2 at the nuclear periphery. Likewise, immunostaining for phosphorylated DDR2 (pDDR2) demonstrated elevated nuclear pDDR2 levels in Aβ-treated cells relative to controls (Fig. 2F, 2G), suggesting that Aβ preferentially promoted nuclear accumulation of DDR2 in its active form.

**Figure 2:**
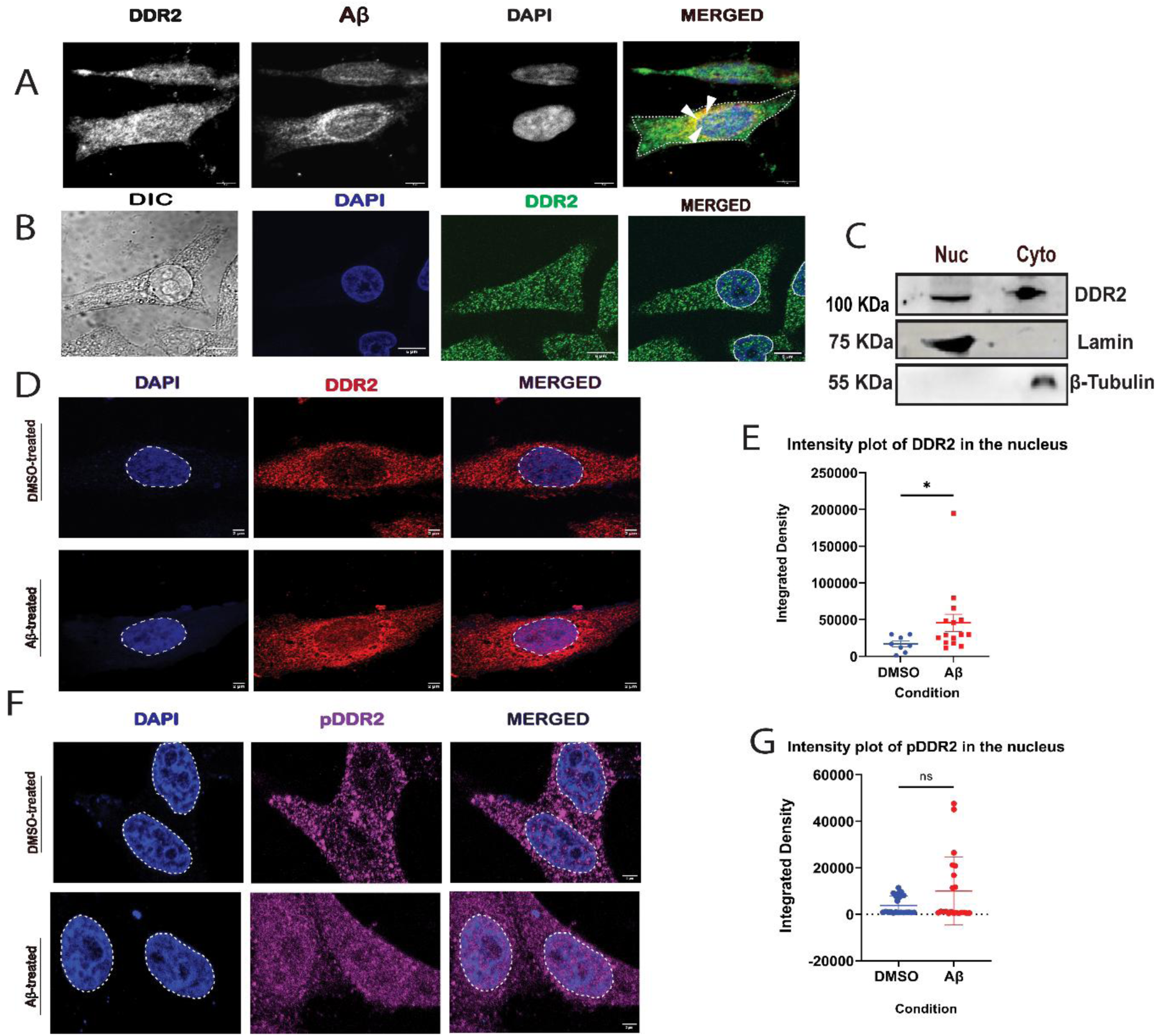
(A) SH-SY5Y cells were transfected with DDR2 and treated with Aβ_42_ for 12 hours, followed by immunostaining with anti-Amyloid-β antibody (red) and anti-DDR2 antibody (green). Representative merged images (red, green and DAPI)) are shown. Colocalization analysis was performed using the Coloc2 plugin in ImageJ, analyzing whole cell. The Pearson correlation coefficient was 0.657 ± 0.029. (B) SH-SY5Y cells were fixed, stained with DAPI (blue), and immunostained with anti-DDR2 antibody to assess nuclear localization. (C) SH-SY5Y cells were fractionated into nuclear and cytoplasmic-enriched fractions, and 50μg of samples from each fraction were analyzed by immunoblotting with the indicated antibodies. β-Tubulin was used as a cytosolic marker and Lamin as nuclear marker. (D) SH-SY5Y cells were treated with Aβ_42_ or DMSO and immunostained with anti-DDR2 antibody; nuclei were counterstained with DAPI. (E) Quantification of nuclear DDR2 intensity was performed by measuring integrated density in DMSO- versus Aβ_42_ treated cells, and the results are presented as a comparative plot. (F) SH-SY5Y cells were treated with Aβ_42_ or DMSO and immunostained with anti-phospho-DDR2 (pDDR2) antibody; nuclei were counterstained with DAPI. (G) Quantification of pDDR2 intensity in DMSO- versus Aβ_42_ treated cells was performed using ImageJ, and the results are presented as a comparative plot.

### WRG-28 inhibits DDR2 activation in a biphasic manner

To check whether Aβ_42_ and collagen shared the same DDR2 binding site and whether the inhibitor WRG-28 could block the Aβ_42_-mediated phosphorylation of DDR2 following a similar mechanism, SH-SY5Y cells were first incubated with Aβ_42_ followed by exposure to increasing concentrations of WRG-28. At a low inhibitor concentration (0.5 µM), a marked increase in the DDR2 phosphorylation was observed. However, with a further increase in inhibitor concentration, the phosphorylation of DDR2 gradually declined demonstrating a dose-dependent inhibitory effect of WRG-28 (Fig 3A). The biphasic plot well depicted the impact of WRG-28 binding on DDR2: Aβ_42_ complex (Fig 3A,iii). This biphasic behaviour suggested that WRG-28 might initially facilitate the activation of DDR2 under certain conditions before inhibiting it at higher concentrations. To put this into perspective we traced WRG-28-mediated inhibition of DDR2 activity triggered by collagen. Unlike before, increasing the concentration of WRG-28 induced a steady deactivation of the kinase indicating different binding sites for the ligands, and their different inhibition mechanism (3B). The whole mechanism has been illustrated in Fig 3C.

**Figure 3:**
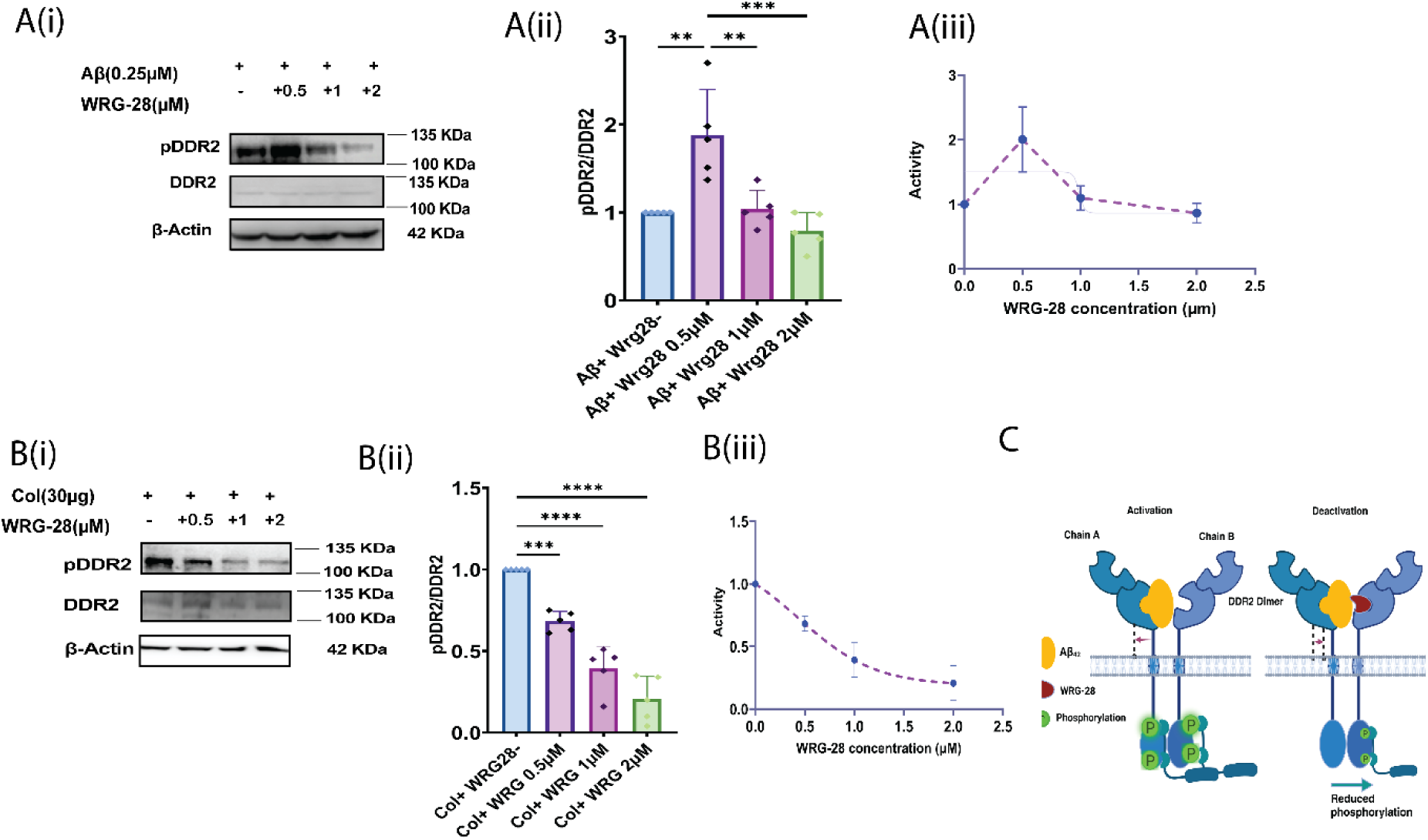
(Ai,ii) SH-SY5Y cells were treated with 0.25 µM Aβ_42_ peptide for 6 hours in the presence of a WRG-28 inhibitor of varying concentration. Cells were harvested, western blotted with pDDR2, and reprobed with DDR2 antibody. Densitometric quantification was performed. A representative blot was shown, and the quantification of three independent experiments was represented as mean ± SEM. (Aiii)The activity of DDR2 was plotted against the inhibitor concentration (0.5µM to 2µM), which represents the biphasic mode of inhibition. (Bi,ii) SH-SY5Y cells were treated with Collagen I for 4 hours in the presence of WRG-28. Cells were collected and subjected to immunoblotting using a pDDR2 antibody to evaluate DDR2 phosphorylation levels. (B iii) The activity of DDR2 was plotted against the inhibitor concentration (0.5µM to 2µM), which represents a steady decline the DDR2 activity (C) The mode of action of WRG-28 binding to DDR2: Aβ_42_ complex has been illustrated sequentially.

### Distinct Binding of Aβ_42_ to DDR2 Independent of the Collagen-Binding Site

Since HEK293 cells express very low levels of endogenous DDR2, it was chosen to show the distinct binding of Aβ_42_. Besides the wild-type clone (DDR2-WT), a single point mutant (E113K) clone (DDR2-Mut) was used, where the collagen-binding site had been mutated. The cells were transfected with either a wild-type DDR2 (DDR2-WT) plasmid or a DDR2 mutant (DDR2-Mut) plasmid and incubated for 24 hours to allow protein expression. Following transfection, cells were treated with either collagen I or amyloid-beta (Aβ_42_) for an additional 6 hours. Subsequently, cells were harvested, lysed, and subjected to immunoblotting to assess DDR2 phosphorylation and downstream signaling effects.

A one-way ANOVA was performed to evaluate the effect of different treatments on DDR2 phosphorylation levels relative to the control group. The results indicated that collagen I treatment increased DDR2 phosphorylation in the wild-type group compared to control cells, confirming that collagen I is an activator of DDR2. Notably, Aβ_42_ treatment resulted in a more significant increase in DDR2 phosphorylation than collagen I treatment. In contrast, DDR2-Mut cells treated with collagen I exhibited a reduced phosphorylation level compared to DDR2-WT + collagen I, suggesting that the mutation disrupts collagen-induced DDR2 activation. However, in DDR2-Mut cells treated with Aβ_42_, phosphorylation levels were significantly higher compared to the DDR2-Mut + collagen I group and were comparable to DDR2-WT + Aβ_42_ (Fig 4).

**Figure 4:**
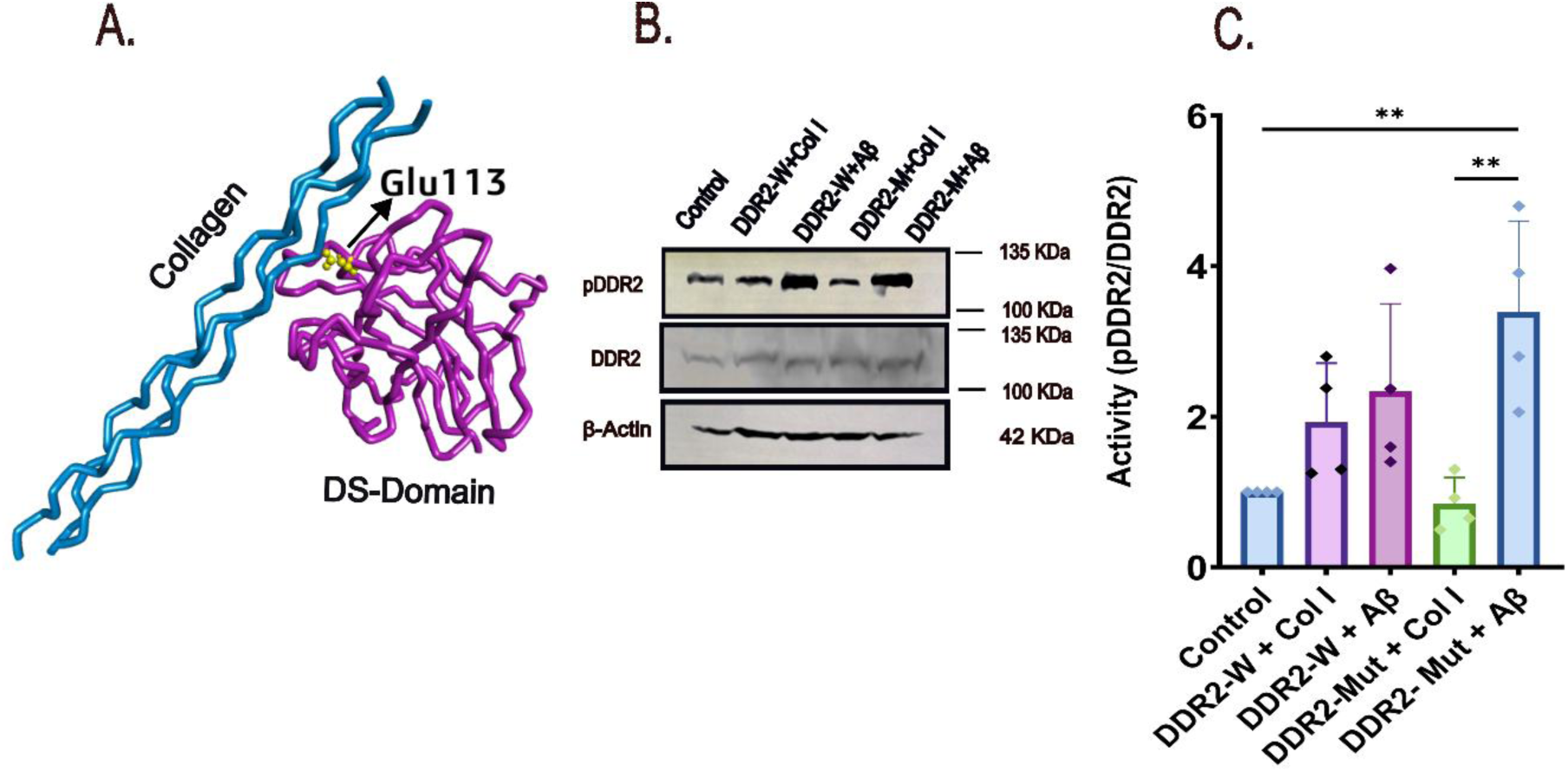
(A) Structural representation of the DDR2-collagen complex highlighting the Glu 113 residue (yellow) with DDR2 (purple) and its proximity to collagen (blue). (B) Immunoblot analysis showing levels of phosphorylated DDR2 (pDDR2), total DDR2 and β-Actin (loading control) across the different treatment groups. (C) Quantification of DDR2 activity under various conditions. Wild-type DDR2(DDR2-WT) and mutant DDR2 (DDR2-Mut) were treated with either collagen or amyloid-beta (Aβ) as indicated. Data are presented as mean ± SEM from four independent experiments.

These findings indicate that Aβ_42_ can induce DDR2 phosphorylation independently of the collagen-binding site. The ability of Aβ_42_ to activate DDR2 in the mutant form supports the hypothesis that Aβ_42_ binds to a site distinct from the collagen I-binding domain, providing further insight into the unique molecular interactions governing DDR2 activation.

### Modeling of DDR2 receptor and ligand docking offers insightful details

Our cellular-level experiments identified Aβ_42_ as a potential ligand for DDR2 in the context of Alzheimer’s disease, with its interaction further inhibited by WRG-28. To understand the mechanistic insight of the entire process at the molecular level, we have simulated a model to show the most probable interaction site of Aβ_42_ binding and WRG-28 interaction. In the absence of the crystal structure of the entire ectodomain, we first predicted the structure up to the TM region using Alphafold2. The DS-domain matched with the available crystal structure (PDB id: 2WUH) and the quaternary disposition of the DS- and DS-like domains resembled its close homolog DDR1 (PDB id: 4AG4). Owing to DDR’s typical prevalence on the cell surface as a dimer, we constructed a dimer model of the DDR2 ectodomain extending up to the TM part (Fig 5A). To validate the models, the DDR1-based and AlphaFold 2-generated, DDR2 structures were superimposed in Discovery Studio, yielding a lower RMSD value, indicating high structural similarity. To decipher the interaction between Aβ_42_ and DDR2 dimer, we performed docking using a pentameric assembly as the oligomeric form of Aβ_42_ is biologically relevant species in Alzheimer’s disease. From the docking studies, the most suitable docking orientation was selected based on the lowest binding energy score in ClusPro (Supplementary Table 2). We have also docked WRG-28 in the presence and absence of Aβ_42_ peptide. Finally, we generate a physiologically relevant membrane to validate the simulated complex. The dimeric assembly has been optimized and all the interaction studies have been performed on the optimized complex structure(s).

**Figure 5:**
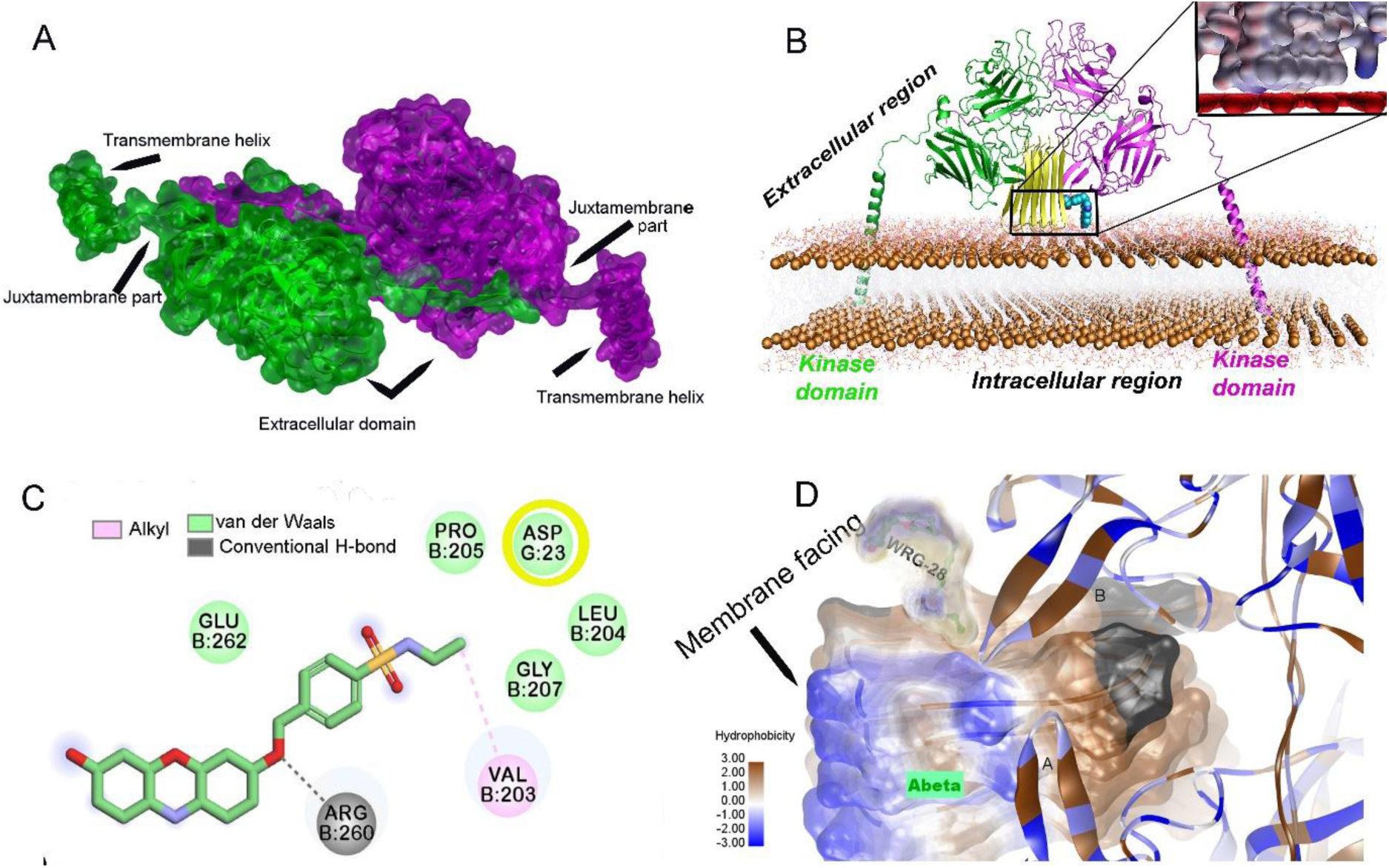
(A) DDR2-dimer generated using ClusPro self-docking is shown perpendicular to the membrane. (B) DDR2-Aβ_42_-WRG-28 complex generated from ClusPro docking positioned inside the membrane bilayer modeled by CHARMM-GUI. Electrostatic interaction exhibited by Aβ_42_ and WRG-28 with the membrane has been shown in the inset. (C) Represents the interactions between WRG-28, the DDR2 receptor and Aβ_42._ Stabilizing non-covalent interaction mediated by the WRG-28 (ball and stick representation) with chain B of DDR2 residues and Van der Waals interaction with G chain of Aβ_42_ peptide (encircled in yellow) in 2D illustration. (D) Aβ_42_ binding to the DS-like domain of DDR2 dimer with the turn region of its ‘β-turn-β’ motif oriented toward the extracellular surface of the membrane bilayer.

DDR2, being a membrane protein, required a suitable environment to maintain its structural integrity and proper orientation, akin to its natural biological setting. To replicate this environment and ensure the accurate positioning of the protein-ligand complex, we utilized CHARMM-GUI to generate a realistic membrane bilayer model. The generated models of DDR2:Aβ_42_:WRG-28 complex were analyzed for their compatibility and the models exhibited favourable electrostatic interactions, aligning well with the membrane surface potential. The fact that the complex could stably associate with the membrane, reflected a physiologically relevant interaction (Fig 5B).

The interaction analysis between WRG-28 and DDR2 receptor revealed that WRG-28 could bind specifically to the DS-like domain of the DDR2, overlapping with the Aβ_42_ binding region. It formed a conventional hydrogen bond with Arg260 and an alkyl interaction with Val203 both part of the DS-like domain. Additionally, a van der Waals interaction was found between WRG-28 and the G-chain of Aβ_42_peptide (Figure 5C). It was further observed that WRG-28 binds in a groove formed by DS-like domain and Aβ_42_peptide, suggesting a stable interaction in presence of Aβ_42_ peptide. This cooperative stabilization is further supported by the improved interaction energy observed in the presence of WRG-28 (Supplementary Table 4) and is illustrated in the membrane-embedded structural context (Figure 5D). Docking results showed that Aβ_42_ was optimally binding to the DS-like domain with its turn region of the ‘β-turn-β’ motif facing the extracellular surface of the membrane bilayer (Figure 5D). This mode of interaction is distinctively different from the collagen-binding site at the DS-domain and it is not surprising as the structural element and composition of Aβ peptides are completely different from triple helical collagen fibres. The interaction between the DDR2 dimer and different chains of Aβ_42_ fibrils has been represented in (Supplementary Figure 3). All the interacting residues have been analyzed using Discovery Studio and are listed in (Supplementary Table 3).

### Aβ_42_ peptide conformationally impacts the Juxtamembrane domain of DDR2 upon binding

When we superimposed the three minimized complexes, DDR2, DDR2:Aβ_42_, and DDR2:Aβ_42_:WRG-28. From the superimposed structures, notable deviations were observed in the juxtamembrane part of the chain to which Aβ_42_ was bound (Chain A in Fig 6A). Specifically, these deviations occurred post-Aβ_42_ binding. The asymmetric binding of Aβ_42_ to DDR2 induced conformational changes within the JM region, in a functionally relevant way. The loop curvature in the JM domain was primarily introduced by Pro378, Pro381, and Pro384. The most deviated amino acids were Pro381 and Met382 with altered rotameric conformations (Fig.6B). This altered conformation of juxtamembrane might be a plausible reason for signal transduction to the intracellular kinase domain. This observation highlights the significance of amyloid-beta binding in modulating the structural configuration of DDR2 (Supplementary Table 5). Interestingly, upon WRG-28 binding to the DDR2:Aβ_42_ complex, a distinct structural shift in the JM region was observed, characterized by a backward displacement. This shift, particularly evident at Pro381, which deviated by 2.694 Å, suggests a conformational reset that counteracts the Aβ_42_-induced activation. The backward displacement of the JM region likely disrupts the signal propagation necessary for kinase activation, leading to the inhibition of DDR2 activity. This finding highlights the potential of WRG-28 as a structural modulator capable of reversing pathological DDR2 activation, offering mechanistic insights into its inhibitory effects (Fig 6C, 6D).

**Figure 6:**
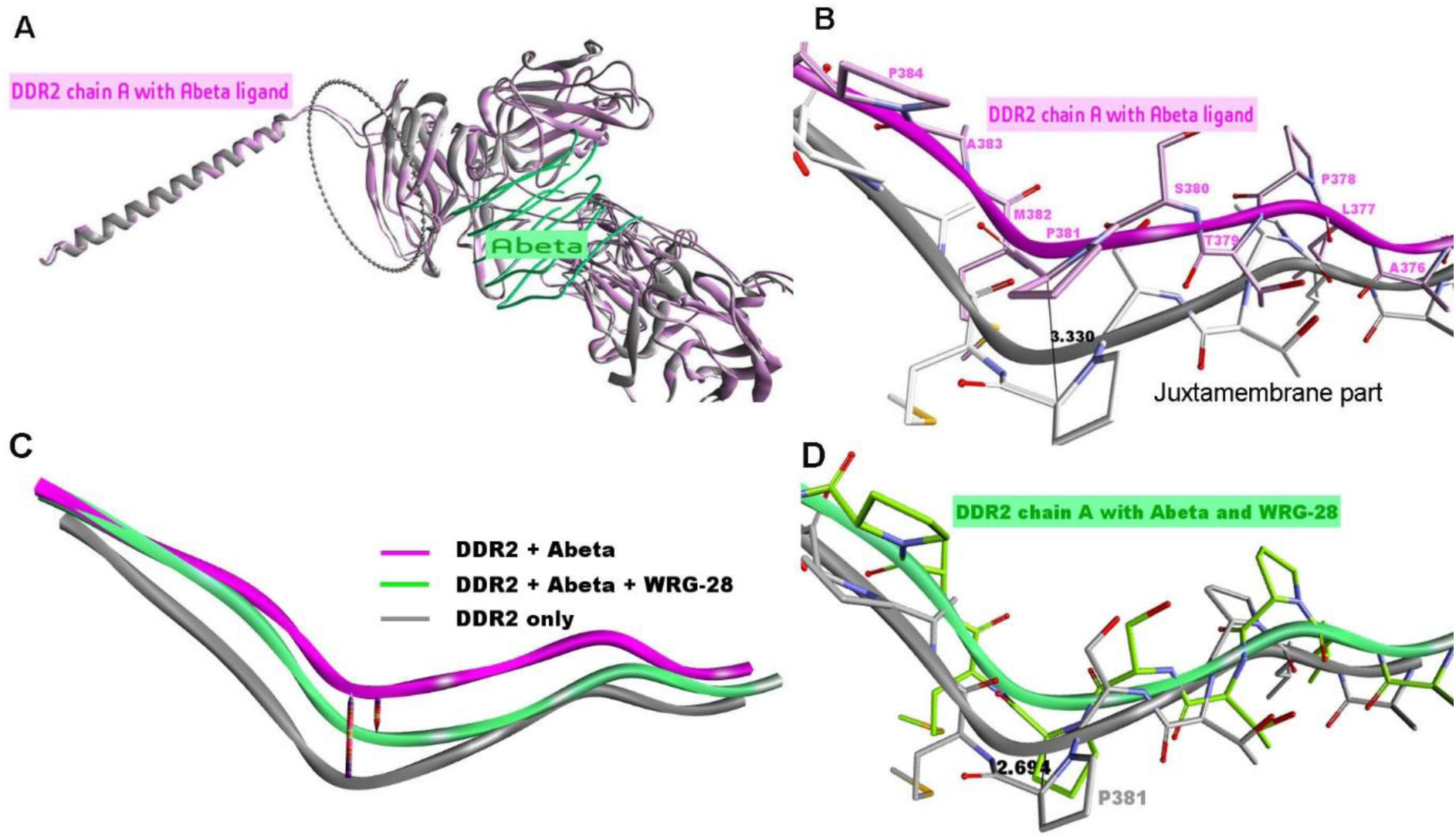
(A) Superimposed structures of the minimized DDR2 complexes highlight the juxtamembrane domain’s structural deviations upon Aβ_42_ binding (represented in pink) (B Close-up view of the DDR2 juxtamembrane (JM) region highlighting the structural deviations induced by Aβ_42_ binding. Pro381 exhibits significant rotameric shifts, with a measured deviation of 3.33 Å. (C) Structural superimposition of DDR2 in its apo form (gray), DDR2:Aβ_42_ complex (purple), and DDR2:Aβ_42_:WRG-28 ternary complex (green) reveals the backward shift in the JM region upon WRG-28 binding. (D) Detailed view of the WRG-28-induced JM displacement. Pro381 undergoes a 2.694 Å shift, indicating a structural reset that likely inhibits DDR2 activation.

After the final energy minimization steps interaction energy was calculated between (1) DDR2: Aβ_42_ and WRG-28, (2) DDR2:WRG-28 and Aβ_42_ (3) DDR2 chain A and DDR2 chain B, (4) DDR2 and Aβ_42_.

Aβ_42_ binding was most favourable when forming a ternary complex with DDR2 and WRG-28 (Supplementary Table 4), primarily driven by electrostatic and hydrogen bonding interactions. Aβ_42_ also exhibited strong binding affinity to DDR2 dimer as well, with electrostatic interactions playing a pivotal role. Specifically, DDR2 residue LYS119 (Chain A) forms multiple electrostatic and hydrogen bond interactions with C-terminal carboxylate groups (OCT1 and OCT2) of Aβ chains C and D. Additionally, GLN139 and TYR211 forms hydrogen bonds with ALA42, GLU22, and ASP23 of Aβ_42_. For WRG-28 binding, however, either it binds to DDR2 first, followed by Aβ_42,_ or vice versa, and either way, its stable binding restricts conformational changes necessary for signal transduction in the kinase domain and would downregulate the kinase activity. Based on the docking scores of Autodock Vina, we excluded the first possibility as the highest docking score was -2.161 Kcal/mol (Supplementary Table 6). Whereas, the docking score of WRG-28 binding to DDR2:Aβ complex was -4.6 Kcal/mol (Supplementary Table 7). So, it can be concluded that WRG makes a ternary complex with DDR2-Aβ and restricts the necessary movement for downstream signal transduction.

## Discussion

This study highlights the dysregulation of Discoidin domain Receptor 2 in Alzheimer’s Disease (AD) pathology, with its activity and expression getting significantly upregulated in AD. Notably, DDR2 can be non-canonically activated by Aβ_42_ even in the absence of its canonical ligand, collagen, suggesting a novel pathological mechanism. This work therefore provides a new perspective on the role of DDR2 and its interaction with Aβ_42_ peptide in Alzheimer’s disease.

Several studies have previously reported that receptor tyrosine kinases can translocate to the nucleus through various mechanisms, both under physiological and pathophysiological conditions^22,24–26^. A notable observation in our study is the nuclear translocation of DDR2 in neuroblastoma cells, both under untreated conditions and after Aβ_42_ treatment. This finding suggests a previously unrecognized functional role for DDR2 in intracellular signaling and gene regulation, potentially linking DDR2 activity to transcriptional changes that contribute to AD pathology. Future studies are warranted to elucidate the downstream effects of DDR2 nuclear localization and its impact on cellular homeostasis in the AD brain

Previously, it has been reported that WRG-28, a known inhibitor of DDR2, binds to the interface of DS and DS-like domain of the receptor and inhibits collagen-I-mediated tyrosine phosphorylation^27^. To further understand the binding site of Aβ_42_ on DDR2, we used WRG-28, and interestingly enough, we observed a biphasic mode of inhibition of activated DDR2: Aβ_42_ complex in the presence of varying concentrations of WRG-28. At lower concentrations (0.5 µM), WRG-28 initially enhances the activity of DDR2, while at higher concentrations (1-2 µM) the activity declines. This paradoxical activation of DDR2 at a lower concentration of inhibitor suggests that WRG-28 may induce structural or conformational changes in DDR2 which facilitate its activation by Aβ_42_. Docking studies also support this sequential binding as WRG-28 is docked more effectively on DDR2 dimer: Aβ_42_ complex, than on the DDR2 dimer alone.

The molecular docking studies revealed that Aβ_42_ binds to the DDR2 dimer at sites distinctly different from the collagen-binding DS domain, indicating a unique binding mechanism independent of collagen interaction. Despite the pseudo-two-fold symmetry of the DDR2 dimer, Aβ_42_ binds asymmetrically owing primarily to the inadequate space inside the binding cavity of the dimer which can accommodate only one Aβ_42_ oligomer at a time (Supplementary Figure 4). Four consecutive peptide chains of the Aβ_42_ interact primarily with one of the chains (chain A) of the dimer and a single peptide chain having contact with the other monomer. Notably, the protruding loop between the β1 and β2 strands of the DS-like domain in the B chain docks into the central groove of a terminal β-turn–β motif of Aβ_42_. In contrast, the corresponding loop of the A chain extends alongside the stacked Aβ peptides, aligned nearly parallel to the fibril axis. The model accommodates continued fibril elongation, supporting the structural feasibility of DDR2–Aβ_42_ interactions during fibril growth. Interestingly in the docking studies using DDR2 dimer, WRG-28 binds to the DS-like domain which is different from the previously reported binding site. When WRG-28 was docked separately on the i) DDR2: Aβ_42_ complex and ii) DDR2 alone, the former showed a higher docking score indicating that probably WRG-28 makes a ternary complex with DDR2-Aβ_42_ initially acting like a wedge that facilitates Aβ_42_ to bind more tightly with DDR2, leading to its activation. Soon WRG-28 binding to its cognate allosteric binding site restricts the JM domain from its necessary movement for downstream signal transduction and hence deactivates the observed unique biphasic activation/ inhibition of DDR2 by WRG-28.

To further validate the unique binding, the site-directed mutagenesis revealed that Aβ_42_ could retain its ability to activate DDR2 while collagen failed as expected in the E113K mutant which is deficient in collagen binding. The ability of Aβ_42_ to independently activate DDR2 in the mutant form supports that Aβ_42_ binds to a site distinct from the collagen I-binding domain, and activates DDR2 in a non-canonical way.

Further, the distinct binding site of Aβ_42_ on DDR2 offers the potential for a targeted therapeutic intervention. Designing inhibitors that specifically block Aβ_42_ binding without affecting collagen binding could mitigate DDR2’s pathological activation in AD while preserving its physiological function.

In conclusion, this study establishes DDR2 as a critical player in AD pathology through its unique activation by Aβ_42_ and subsequent nuclear translocation in the active form. It further highlights the complex interplay between DDR2-Aβ_42_-WRG-28. It paves the way for future investigation of DDR2’s role in AD and downstream signaling effects resulting from amyloid-beta binding, with the ultimate goal of developing DDR2-targeted therapies that could alleviate disease progression and improve patient outcomes.

## Materials and Methods

### Cell culture and transfection

The Human Neuroblastoma cell line (SH-SY5Y) was procured from the official repository of the National Centre for Cell Sciences, Pune, India. The SH-SY5Y cells were cultured following the manufacturer’s protocol in DMEM-F12 media (Gibco), supplemented with 10% fetal bovine serum (FBS) (Gibco) under humified condition at 37^◦^C and 5% CO_2._ The transfection of the cells was all carried out using the transfection reagent Lipofectamine 2000 (Invitrogen) with cells reaching 70-80% confluency. After 48 hours of transfection, the cells were monitored for their transfection efficiency by observing the GFP expression under a fluorescence microscope and were used further for experiments.

HeLa (Human cervical cancer line) and HEK 293 (Human embryonic kidney 293) were a gift from Prof. Oishee Chakraborty (Saha Institute of Nuclear Physics). Both the cells were cultured using the manufacturer’s protocol in Dulbecco’s modified Eagle’s medium (DMEM; Himedia) supplemented with 10% fetal bovine serum (FBS; Himedia) at 37^◦^ and 5% CO_2_. The transfection of cells was carried out using Lipofectamine 2000 (Invitrogen). After 48 hours of transfection, cells were monitored and used for experimental purposes.

### Plasmid constructs, Reagents, and antibodies

The PGFP-C1 (Clontech) (referred to as GFP in the text) and GFP-tagged AICD (APP intracellular domain) construct were already available in the lab ^28,29,30,31,32^. Reagents: WRG-28 (SML2786), Rat Collagen type I (C7661). DDR2 antibody (ab63337) pDDR2 antibody (MAB25382) Anti-Lamin antibody ( ab 190380), HRP-anti-beta Actin antibody( mAbcam 8226). The primers used in this study are listed in Supplementary Table 1.

### AD Cell Model

SH-SY5Y cells were transfected with either AICD-GFP or GFP (control) using the Lipofectamine 2000 transfection reagent. The lyophilized Aβ_42_ protein fragment (Sigma, A980) was dissolved in DMSO to create a 2 mM stock solution. Three hours after transfection, GFP-transfected cells were treated with DMSO, while AICD-transfected cells were treated with 0.5 μM Aβ_42_. The samples were collected 48 hours after treatment. GFP-transfected cells treated with DMSO served as the control, while AICD-transfected cells treated with Aβ_42_ represented the Alzheimer’s disease (AD) cell model^33,34,35^ (Fig 1B).

### RNA Isolation, cDNA Conversion, and Quantitative Real-time PCR

Following the manufacturer’s instruction, total RNA was isolated from cells using TRIzol reagent (Invitrogen) and was quantified using a Nanodrop 2000 microvolume spectrophotometer (Thermo Scientific). 2μg of total RNA was used for each reaction for cDNA conversion using oligo dT as primers for coding genes. The cDNA conversion was done manually using 5 x reaction buffer (Thermo Scientific), 10mM dNTP mix (Thermo Scientific), Oligo dT/random hexamers, and reverse transcriptase enzyme (Thermo Scientific). The qRT-PCR was performed using SYBR green 2X universal PCR Master Mix (Applied Biosystems) in QuantStudio^TM^ 3 Real-Time PCR system (Applied Biosystems). The fold change values of the target genes were calculated using the 2^-ΔΔct method after normalizing with the housekeeping gene.

### Subcellular Fractionation

Cells were washed in ice-cold PBS and harvested in 500ul of Hypotonic Lysis Buffer (HLB) containing (250mM Sucrose, 20mM HEPES-KOH, pH 7.4, 10mM KCl, 150mM MgCl_2,_ 1mM EDTA, 1mM EGTA, 0.5mM PMSF) by scraping and incubated on ice for 15 mins. Using a 1ml syringe the cell suspension was passed through a 27 gauge needle several times until all cells were lysed. It was then kept on ice for 20 mins and centrifuged at 720g for 5 mins. The pellet contains the nuclei and the supernatant contains the cytosol. The nuclear pellet was washed thrice with 500ul HLB and lysed in 5X sample buffer. Both the fractions were run on 8% SDS PAGE gel for immunoblotting.

### Western Blot

Cells were washed three times in ice-cold Phosphate buffer saline (PBS) and lysed on ice in lysis buffer (1M Tris-HCl, pH 7.5, 1 N NaCl, 0.5 M EDTA, 1 M NaF, 1M Na_3_VO_4_, 10% SDS, 20mM PMSF, 10% TritonX-100, 50% glycerol) for 30 min in the presence protease inhibitor (Roche Diagnostics) and centrifuged for 15mins at 13,000 g and supernatants were collected. Protein estimation was performed using Bradford spectrophotometric assay.

Samples were mixed with 5X Laemmli buffer, boiled at 95◦C, and, separated on an 8-12% SDS-PAGE according to their molecular weight. Proteins were then transferred on a PVDF membrane (Millipore Corporation) which was blocked with 5% BSA in TBST (50mM Tris-HCl, 150mM NaCl, pH 7.5, 0.05% Tween 20). After transfer, the membrane was probed with primary antibody followed by incubation with HRP-conjugated secondary antibody. The membranes were then developed using an ECL kit (Pierce^TM^) in Chemidoc (Azure Biosystems). Band intensities were measured using GelQuant Express analysis software (DNR Bio-Imaging Systems LTD). All the experiments were repeated thrice or more and band intensities were normalized to loading control.

### Immunocytochemistry

Immunocytochemistry was performed on fixed cells. At first, cells were fixed using 4% paraformaldehyde in PBS for 10 mins at room temperature in the dark. Afterward, cells were washed three times in ice-cold PBS and permeabilized using 0.1-0.25% Triton X-100 for 5-7 mins at room temperature followed by ice-cool PBS wash thrice. Cells were then blocked with 1% BSA in PBS for 2 hours at room temperature to block unspecific antibodies. Then, cells were incubated with primary antibody diluted in 1% BSA in PBS in a humified chamber overnight at 4◦C. After that, the primary antibody was decanted and washed in ice-cold PBS thrice for 5 mins. Cells were then incubated with a secondary antibody diluted in 1% BSA in PBS and kept in the dark for 2 hours, following which the secondary antibody was drained out and washed in ice-cold PBS (thrice for 5 mins). Finally, the cells were incubated with 0.1-1 ug/ml DAPI (DNA stain) for 5mins. The DAPI solution was discarded and rinsed twice with PBS. Coverslips were mounted on previously cleaned and dried slides with a drop of mounting media and sealed with nail polish to prevent drying and movement under the microscope.

### WRG-28 binding assay

SH-SY5Y cells were seeded on a 6-well plate and were treated with Aβ_42_ (0.25 µM) first followed by WRG-28 treatment sequentially with increasing concentration (0.5,1,2 µM). For the control set none of these were added. 6 hours following treatment, cells were harvested and immunoblotted with DDR2 (1:500) and pDDR2 (1:20000) antibodies to assess the activity.

### Generation of DDR2 Mutants

The Full-length cDNA ORF clone of DDR2 transcript variant 1 cloned into the expression vector pCMV3-C-GFP was obtained from Sino Biological. E113K^36^ mutation was generated using GeneArt™ Site-Directed Mutagenesis PLUS Kit and further verified by DNA Sequencing. The primers for the site-directed mutagenesis to introduce missense mutations were as follows:

DDR2-E113K-F: GAGGTCATGGCATC**AAG**TTTGCCCCCATG;

DDR2-E113K-R: CATGGGGGCAAA**CTT**GATGCCATGACCTC.

HEK293T cells were transfected with the wild-type DDR2 plasmid and DDR2 mutant.

### *In-silico* Structure Studies, Docking and Energy Minimization

The DDR2 protein sequence was obtained from the NCBI database. Following its retrieval, a BLASTp (Basic Local Alignment Search Tool for proteins) analysis was conducted to identify homologs of DDR2. DDR1 exhibited 97% query coverage and 54.88% identity. The crystal structure of the DDR1 extracellular region, available in the PDB (PDB ID: 4AG4) at a resolution of 2.80 Å, was used as a template for homology modeling. Using SWISS-MODEL (https://swissmodel.expasy.org/), the extracellular region of DDR2 was modeled. However, since this model did not include the juxtamembrane and transmembrane domains, AlphaFold 2(https://colab.research.google.com/github/sokrypton/ColabFold/blob/main/AlphaFold2.ipynb) was employed to generate a complete DDR2 structure comprising residues 1-430, covering both the extracellular and transmembrane domains. The Aβ_42_ peptide structure was retrieved from the PDB (PDB ID: 2BEG), a well-characterized solid state NMR model. Although missing the flexible N-terminal residues the 2BEG structure retains the key hydrophobic domains (residues 17-21, 30-42) crucial for protofibril formation and docking studies. Its regular β-sheet architecture supports the construction of stable oligomers like pentamers and provide a reliable model for protein-peptide interaction studies.

The structure of WRG-28, a DDR2 inhibitor, was generated from its SMILES code using the NovoPro online tool which generated a 3D structure of the inhibitor (https://www.novoprolabs.com/tools/smiles2pdb).

The ClusPro 2.0 (https://cluspro.org/home.php) protein-protein docking algorithm was used for docking purposes. This algorithm works in three steps. Firstly, for rigid body docking it runs PIPER, based on a Fast Fourier transform (FFT) approach; secondly, it uses a clustering approach to find highly populated clusters of low-energy conformations and discards the unstable clusters. Finally, it runs CHARMM minimization to remove steric clashes. We chose our best-fitted model based on the cluster size and parameters generated by balanced, electrostatic, hydrophobic and Van der Waals interactions. We resorted to two different kinds of docking protocols: (1) the manually optimized monomer structure of DDR2 generated from AlphaFold2 to generate DDR2 dimer using protein-protein docking, (2) Aβ_42_ (PDB:2BEG) and DDR2 docking. In absence of a full-length high resolution DDR2 structure containing transmembrane and cytosolic domains, our model represents a functional approximation. The transmembrane domains were included solely to maintain topological context and were not optimized to reproduce leucine zipper mediated helix-helix interaction^37^. Their observed spatial separation likely reflects membrane embedding constraints and the limitations of CHARMM based energy minimization, which was not tailored to enforce transmembrane domain interactions. This approximation was deemed sufficient for the evaluation of ligand receptor interaction in the extracellular domain which was the primary focus of our docking analysis.

The *in-silico* docking of WRG-28 on the DDR2 dimer was done using AutoDock Vina. A grid box with dimensions of 20 Å × 20 Å × 20 Å was generated, centered at coordinates -19 Å × 17 Å × 37 Å. All the top 20 generated models were clustered around the same binding site The top model, with an affinity score, was selected for further analysis. Next, DDR2 being a membrane protein, it is imperative to know the exact orientation of the molecule concerning the membrane bilayer and the exact position of the transmembrane region hence, a membrane bilayer was generated and the 3D structure of the protein-ligand complex was embedded inside the lipid bilayer to match its native conformation before running the energy minimization. The CHARMM-GUI (https://www.charmm-gui.org/) Membrane/ Bilayer builder module ^38,39,40,41^. The 3D coordinates of the modelled DDR2 dimer docked with Aβ_42_ oligomer and WRG-28 were uploaded to CHARMM-GUI, and the Bilayer Membrane Builder module was used to incorporate membrane components into the system. Before membrane construction, careful orientation and proper positioning of the complex were performed, with the structures oriented at angles of X = 25◦, Y = 142◦ Z = -65 Å. Additionally, the molecule was flipped along the Z-axis to ensure proper alignment with the membrane. The system size was set to 200 Å, and the lipid type POPE (phosphatidylethanolamine) was selected, maintaining a lipid ratio of 1:1. To simulate a realistic biological environment, water molecules and ions were added to the system. The entire setup, including the membrane, water, and ions, was subsequently assembled and equilibrated to prepare the model for downstream simulations and analyses. This step ensured that the model closely resembles the physiological conditions under which the DDR2 dimer and Aβ peptide interact.

The entire DDR2: Aβ_42_: WRG-28 complex underwent an initial energy minimization to resolve steric clashes and optimize binding interactions. Now from that minimized structure, three different complexes were extracted: (1) DDR2 dimer (2) DDR2:Aβ_42_ (3) DDR2: Aβ_42_:WRG-28. All these minimized complexes were solvated independently in an orthorhombic box containing TIP3P water molecules with 7Å minimum distance between the protein atoms and the edge of the boundary box. Na^+^ and Cl^-^ were added to the system to maintain charge neutrality. The solvated structures were first minimized using 500 cycles of energy minimization using CHARMM forcefield with harmonic restraints applied to the protein backbone, allowing side-chain relaxation. This was followed by 2000 additional minimization cycles using the smart minimizer in Discovery Studio (https://www.3ds.com/products/biovia/discovery-studio), during which harmonic constraints were applied specifically to the backbone atoms of the transmembrane helices. This approach approximates membrane constraints and mimics the stabilizing effect of the lipid bilayer, enabling us to observe relevant conformational changes, particularly in the juxtamembrane region. While long-timescale molecular dynamics (MD) simulations were not performed due to computational limitations, this constrained minimization strategy allowed us to capture functionally meaningful structural deviations that may be linked to DDR2 signal transduction. Interaction energies were calculated separately for all three minimized complexes using BIOVIA Discovery Studio.

## Authors Contribution

**RD**: Conceptualization, Original Draft, Methodology, & Data Analysis; **SB**: Conceptualization, Methodology, & Supervision; **DM**: Funding Acquisition, Original Draft editing, & Supervision.

## Data availability

All data presented in this manuscript are included within the document.

## Notes

All the illustrations of this paper were created using BioRender (https://app.biorender.com/biorender-templates/figures).

## Acknowledgments

We thank Prof. Oishee Chakraborty for providing us with HEK and HeLa cells. RD would like to acknowledge UGC for her fellowship.

## Abbreviations

DDR2: Discoidin Domain Receptor 2
Aβ_42_: Amyloid beta 42
RTK: Receptor Tyrosine Kinase

