## Supplementary Figure 1 for "Aβ_42_ Facilitates the Activation of Discoidin Domain Receptor 2 and its Nuclear Enrichment in Alzheimer’s Disease Model"

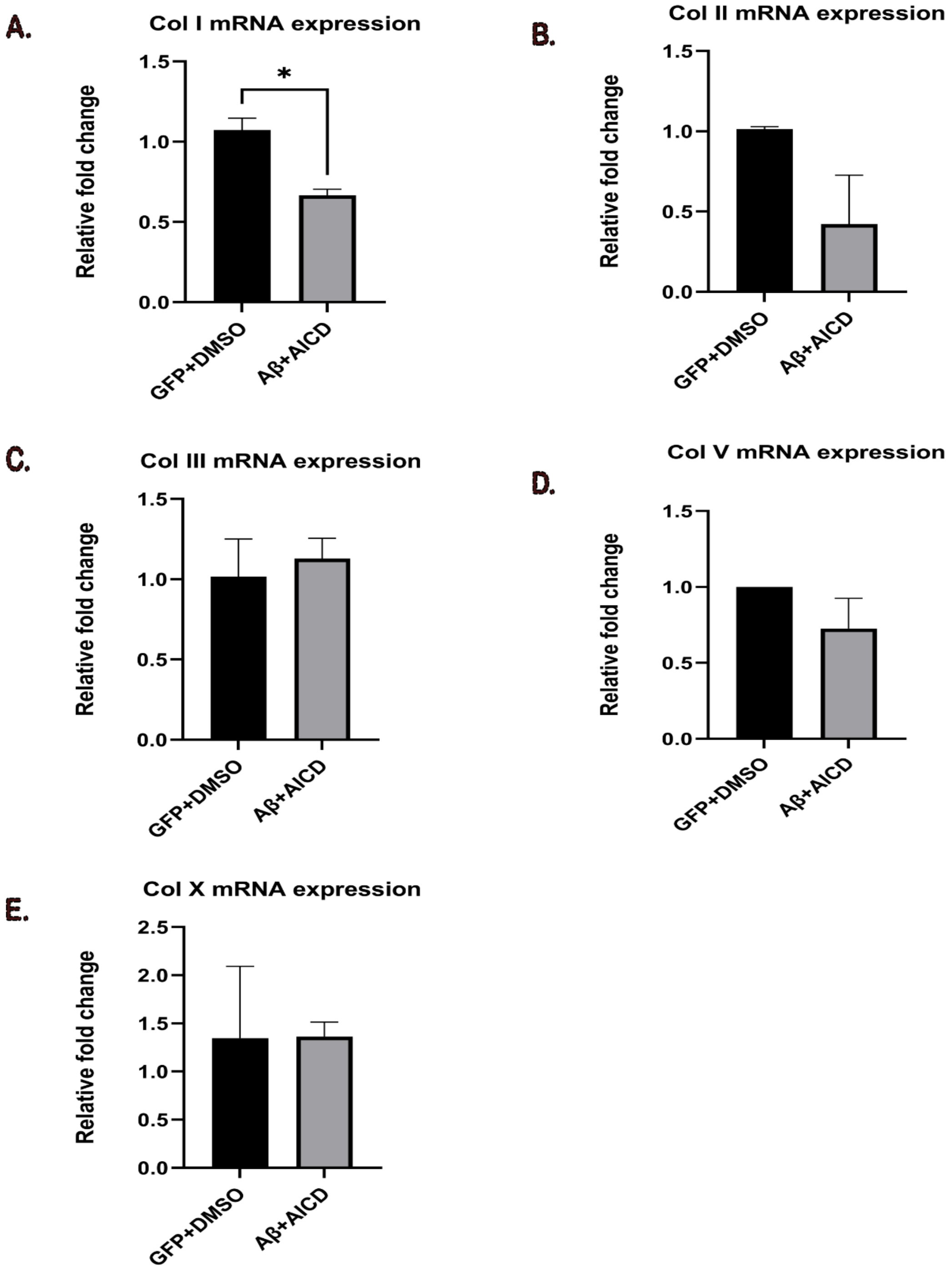


**Supplementary Figure 1**: **The relative fold change of collagen (Col 1, Col 2, Col 3, Col 5, Col 10) transcript level was analyzed in AD cell (Aβ+AICD) compared to control (GFP+DMSO). Col 1 was significantly downregulated in AD (-0.4067 ± 0.08239) compared to control. Col 2 was non-significantly downregulated in AD (-0.5929 ± 0.2157). There was no significant difference in Col 3 expression (0.1129 ± 0.1531). Col 5 was slightly downregulated but was non-significant (-0.2726 ± 0.1984). For Col 10 there were significant changes observed (0.01605 ± 0.7601).**
