## Supplementary Figure 2 for "Aβ_42_ Facilitates the Activation of Discoidin Domain Receptor 2 and its Nuclear Enrichment in Alzheimer’s Disease Model"

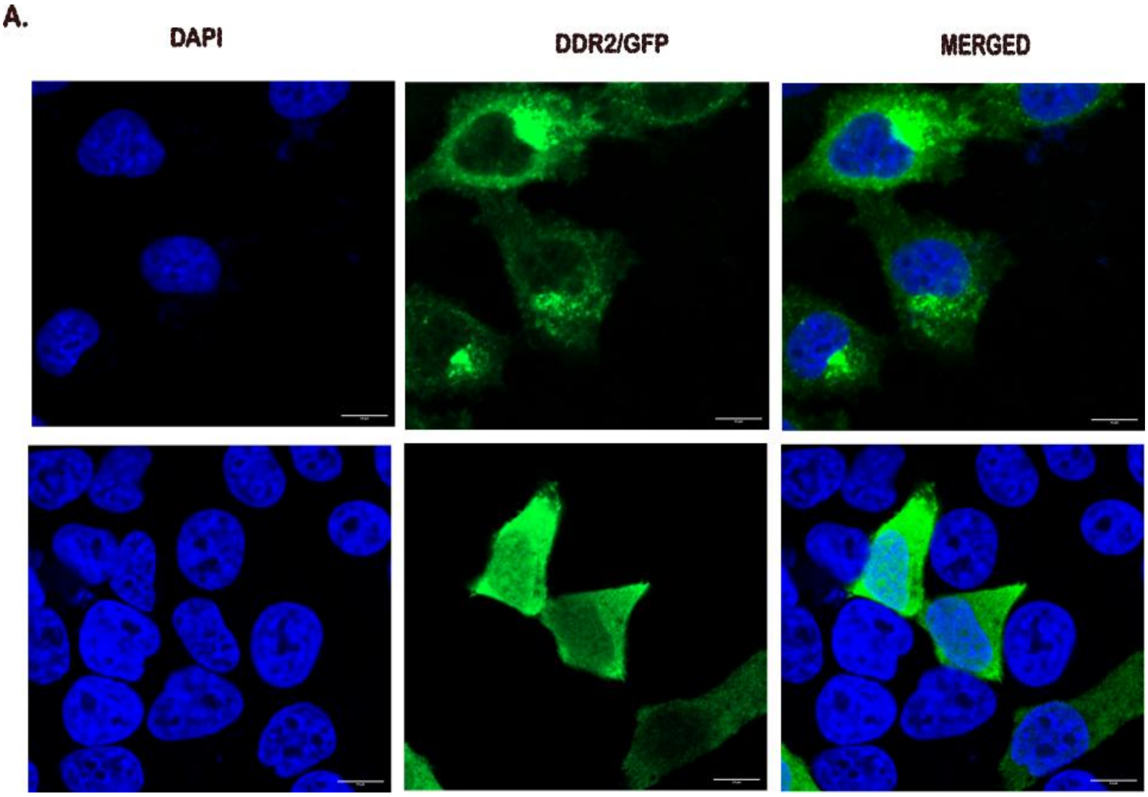


**Supplementary Figure 2**: GFP-tagged DDR2 was transfected in Hela cells and was immunostained with anti-DDR2 antibody to assess the nuclear localization of DDR2
