## Supplementary Figure 3 for "Aβ_42_ Facilitates the Activation of Discoidin Domain Receptor 2 and its Nuclear Enrichment in Alzheimer’s Disease Model"

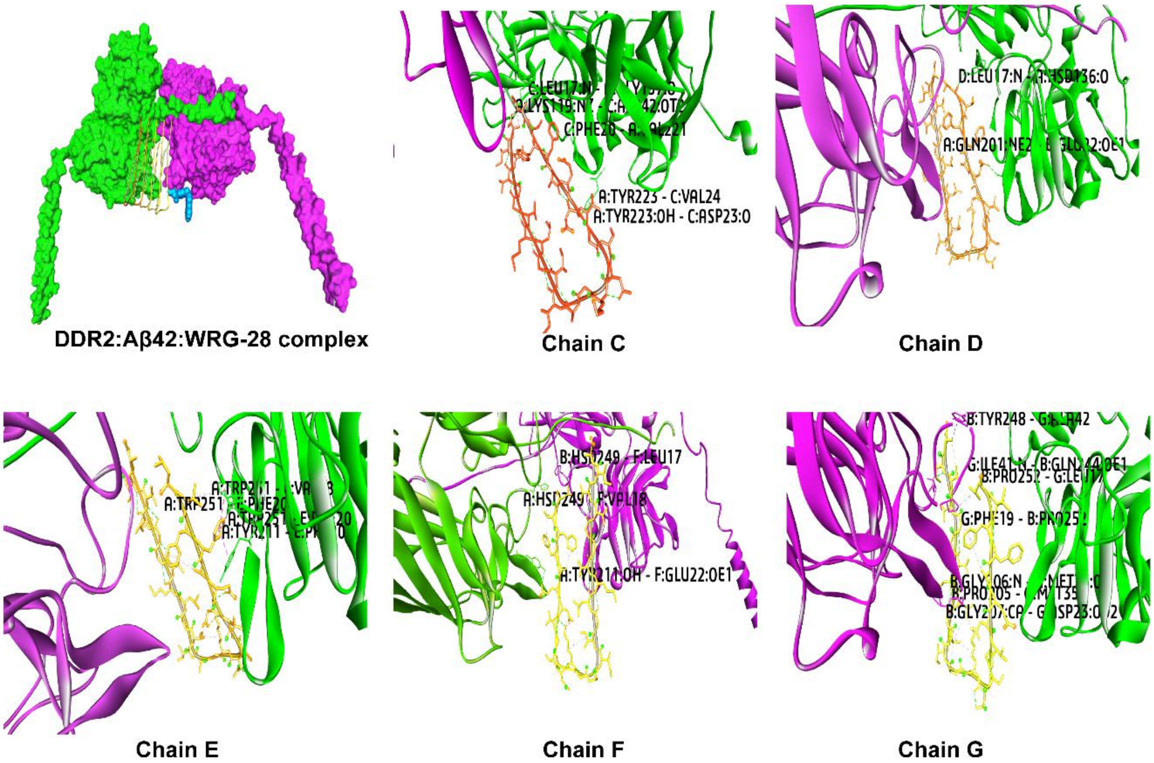


**Supplementary Figure 3**: Represent the interacting amino acid residues of DDR2 and different chains of Aβ42 separately
