## Supplementary Figure 4 for "Aβ_42_ Facilitates the Activation of Discoidin Domain Receptor 2 and its Nuclear Enrichment in Alzheimer’s Disease Model"

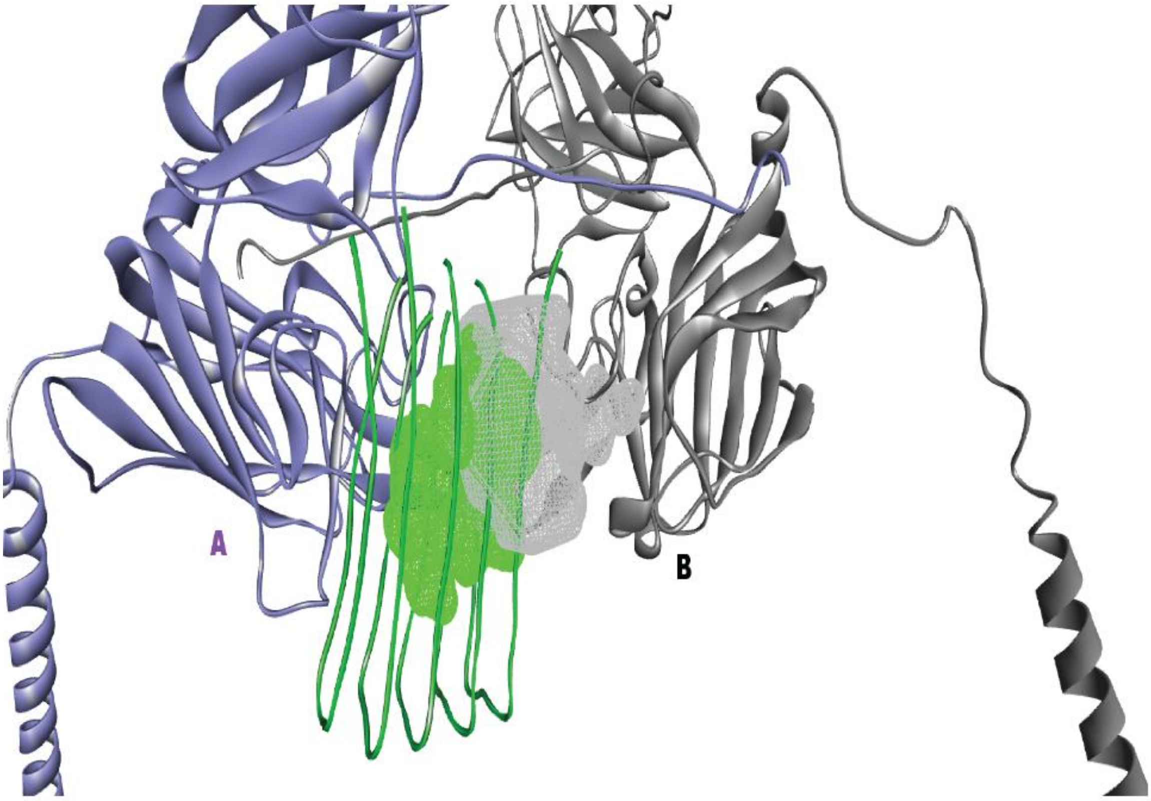


**Supplementary Figure 4: Asymmetric binding of A$\beta_{42}$ to DDR2 dimer.**

This figure illustrates the specific interaction between A$\beta_{42}$ and the DDR2 dimer, highlighting its asymmetric binding mode. The DDR2 binding groove is represented in grey, while the Aβ42 surface is depicted in green. Notably, only a small portion of A$\beta_{42}$ overlaps with the DDR2 binding groove, while the remaining part extends along chain A of the DDR2 dimer. This binding orientation suggests that A$\beta_{42}$ preferentially interacts with chain A, potentially restricting the binding of an additional A$\beta_{42}$ molecule at the same site, thereby allowing only one A$\beta_{42}$ to engage with DDR2 at a time.
