## Supplementary Table 1 for "Aβ_42_ Facilitates the Activation of Discoidin Domain Receptor 2 and its Nuclear Enrichment in Alzheimer’s Disease Model"

**Supplementary Table 1: List of Primers**

| Oligo Name | Sequence 5' to 3' |
|---|---|
| COL1A1_F | TCTGCGACAACGGCAAGGTG |
| COL1A1_R | GACGCCGGTGGTTTCTTGGT |
| COL2A1_F | CTTCACTGGTCTGCAGGGTC |
| COL2A1_R | GGCCAGGATTCCATTAGCA |
| COL3A1_F | CCTGAAGCTGATGGGGTCAA |
| COL3A1_R | TAGTCTCACAGCCTTGCGTG |
| COL5A1_F | TGCATTTCCCGAGGACTTCT |
| COL5A1_R | GTAGAGGAAGACGGGAGAGC |
| COL10A1_F | AAGGGAGAAAGAGGACCTGC |
| COL10A1_R | TGGCCCTGTCTCACCTTTAG |
| DDR2_F | GGAGGTCATGGCATCGAGTT |
| DDR2_R | GAGTGCCATCCCGACTGTAATT |
