## Supplementary Table 2 for "Aβ_42_ Facilitates the Activation of Discoidin Domain Receptor 2 and its Nuclear Enrichment in Alzheimer’s Disease Model"

**Docking scores of DDR2 dimer and A$\beta_{42}$**

Docking scores of DDR2 dimer and A$\beta_{42}$ docking using ClusPro. The table presents the **most densely populated clusters**, representing the most probable binding conformations, along with their respective **cluster centers**. Additionally, the **lowest energy scores** for each docking pose are included, indicating the most thermodynamically favorable interactions. These docking parameters were used to determine the optimal binding orientation of A$\beta_{42}$ with DDR2, which was further analyzed for structural and functional insights.

| Cluster | Members | Representative | Weighted Score |
|---|---|---|---|
| 0 | 91 | Center | -1268 |
| 0 | 91 | Lowest Energy | -1268 |
| 1 | 66 | Center | -1064.1 |
| 1 | 66 | Lowest Energy | -1195 |
| 2 | 60 | Center | -1061.3 |
| 2 | 60 | Lowest Energy | -1190.8 |
| 3 | 42 | Center | -1143.3 |
| 3 | 42 | Lowest Energy | -1182.9 |
| 4 | 38 | Center | -1059.8 |
| 4 | 38 | Lowest Energy | -1240.4 |
| 5 | 38 | Center | -1059.4 |
| 5 | 38 | Lowest Energy | -1195 |
| 6 | 34 | Center | -1070.4 |
| 6 | 34 | Lowest Energy | -1156.6 |
| 7 | 30 | Center | -1260.9 |
| 7 | 30 | Lowest Energy | -1260.9 |
| 8 | 28 | Center | -1060.6 |
| 8 | 28 | Lowest Energy | -1257.4 |
| 9 | 27 | Center | -1088.7 |
