## Supplementary Table 3 for "Aβ_42_ Facilitates the Activation of Discoidin Domain Receptor 2 and its Nuclear Enrichment in Alzheimer’s Disease Model"

**Interacting residues between DDR2 and A$\beta_{42}$**

Molecular Interactions Between DDR2 (A & B) and Amyloid Beta (C–G) are listed in this table and were analyzed using Discovery Studio. The table details key interaction types, including **hydrogen bonds, hydrophobic interactions, electrostatic forces, and van der Waals contacts**, which contribute to the stability and specificity of the DDR2–A$\beta_{42}$ complex.

| DDR2 (Chains A & B) | Residue | Amyloid Beta (Chains C–G) | Residue | Interaction Type |
|---|---|---|---|---|
| A | LYS119:NZ | C | ALA42:OCT2 | Electrostatic |
| A | LYS119:NZ | D | ALA42:OCT2 | Electrostatic |
| A | LYS119:NZ | D | ALA42:OCT1 | Hydrogen Bond |
| A | GLN139:NE2 | C | ALA42:OCT1 | Hydrogen Bond |
| A | VAL203:N | F | GLU22:OE2 | Hydrogen Bond |
| A | TYR211:OH | F | GLU22:OE1 | Hydrogen Bond |
| A | TYR223:OH | C | ASP23:O | Hydrogen Bond |
| B | GLY207:N | G | ASP23:OD2 | Hydrogen Bond |
| B | TYR248:OH | G | ALA42:OCT2 | Hydrogen Bond |
| B | LYS294:NZ | G | GLY37:O | Hydrogen Bond |
| A | LYS138:CA | E | ALA42:OCT1 | Hydrogen Bond |
| B | HSD249:CE1 | F | ALA42:OCT1 | Hydrogen Bond |
| A | ILE209:N | G | PHE20 | Hydrogen Bond |
| A | HSD136:O | D | LEU17:N | Hydrogen Bond |
| A | GLY137:O | D | LEU17:N | Hydrogen Bond |
| B | HSD249:ND1 | F | LEU17:N | Hydrogen Bond |
| A | TYR211:OH | F | ALA21:N | Hydrogen Bond |
| B | HSD249:O | G | LEU17:N | Hydrogen Bond |
| B | VAL250:O | G | LEU17:N | Hydrogen Bond |
| A | VAL221:CG1 | C | PHE20 | Hydrophobic |
| B | PRO205 | G | LEU34 | Hydrophobic |
| A | LYS138 | E | ALA42 | Hydrophobic |
| A | TRP251 | E | VAL18 | Hydrophobic |
| B | TYR248 | G | ALA42 | Hydrophobic |
| B | HSD249 | F | ALA42 | Hydrophobic |
| A | PRO252 | G | PHE20 | Hydrophobic |
