## Supplementary Table 4 for "Aβ_42_ Facilitates the Activation of Discoidin Domain Receptor 2 and its Nuclear Enrichment in Alzheimer’s Disease Model"

**Calculation of Interaction energy**

After energy minimization interaction energy was calculated between (1) DDR2: A$\beta_{42}$ and WRG-28, (2) DDR2:WRG-28 and A$\beta_{42}$ (3) DDR2 chain A and DDR2 chain B, (4) DDR2 and A$\beta_{42}$.

| **Complex** | **Interaction Energy(kcal/mol)** | **Interaction Energy (kcal/mol)** | |
|---|---|---|---|
| | | van der Waals | Electrostatic |
| DDR2-Aβ + WRG | -63.311 | -11.405 | -51.905 |
| DDR2-WRG+ Aβ | -748.48 | -155.762 | -592.723 |
| DDR2-A + DDR2-B | -414.523 | -180.271 | -234.251 |
| DDR2+ Aβ | -591.77 | -169.324 | -422.454 |
