## Supplementary Table 5 for "Aβ_42_ Facilitates the Activation of Discoidin Domain Receptor 2 and its Nuclear Enrichment in Alzheimer’s Disease Model"

**Deviation of the amino acid residues in the juxtamembrane domain**

Structural deviations in the juxtamembrane domain following energy minimization of DDR2 complexes. After minimization, the three complexes— (1) DDR2 dimer, (2) DDR2:Aβ42, and (3) DDR2:Aβ42:WRG-28—were superimposed using Discovery Studio. This table reports the observed deviations in the juxtamembrane region between the DDR2 dimer and DDR2:Aβ42 complex, highlighting conformational changes induced by Aβ42 binding. These structural alterations may provide insights into the activation mechanism of DDR2 and its modulation by amyloid beta

| Residue No. | Residue Name (3 letter) | RMSD value |
|---|---|---|
| 374 | SER | 0.471 |
| 375 | GLU | 1.387 |
| 376 | ALA | 1.306 |
| 377 | LEU | 1.834 |
| 378 | PRO | 2.507 |
| 379 | THR | 2.463 |
| 380 | SER | 2.746 |
| 381 | PRO | 3.335 |
| 382 | MET | 3.378 |
| 383 | ALA | 2.845 |
| 384 | PRO | 1.651 |
| 385 | THR | 1.410 |
| 386 | THR | 2.118 |
| 387 | TYR | 1.928 |
| 388 | ASP | 1.796 |
| 389 | PRO | 1.532 |
| 390 | MET | 1.691 |
| 391 | LEU | 1.701 |
| 392 | LYS | 1.070 |
