## Supplementary Table 6 for "Aβ_42_ Facilitates the Activation of Discoidin Domain Receptor 2 and its Nuclear Enrichment in Alzheimer’s Disease Model"

**Docking scores of DDR2 and WRG-28**

Docking scores of DDR2 and WRG-28 using AutoDock vina. The table represents the binding affinity scores in (Kcal/mol) indicating, the strength and stability of the DDR2-WRG-28 interaction. Lower (more negative) docking scores represent stronger binding and greater thermodynamic favourability. Additionally, the table includes details on the most favourable binding poses, ranked based on their docking scores

| MODEL | ENERGY (Kcal/mol) |
|---|---|
| Model 1 | -2.242 |
| Model 2 | -2.166 |
| Model 3 | -2.133 |
| Model 4 | -2.094 |
| Model 5 | -1.979 |
| Model 6 | -1.973 |
| Model 7 | -1.964 |
| Model 8 | -1.918 |
| Model 9 | -1.852 |
| Model 10 | -1.800 |
