## Supplementary Table 7 for "Aβ_42_ Facilitates the Activation of Discoidin Domain Receptor 2 and its Nuclear Enrichment in Alzheimer’s Disease Model"

**Docking scores of DDR2- Aβ$_{42}$ complex and WRG-28**

Docking scores of DDR2- Aβ42 complex and WRG-28 using AutoDock vina. This table shows that WRG-28 binds more effectively to the DDR2- Aβ42 complex.

| MODEL | ENERGY (Kcal/mol) |
|---|---|
| Model 1 | -4.151 |
| Model 2 | -4.143 |
| Model 3 | -3.839 |
| Model 4 | -3.830 |
| Model 5 | -3.614 |
| Model 6 | -3.569 |
| Model 7 | -3.521 |
| Model 8 | -3.398 |
| Model 9 | -3.319 |
| Model 10 | -3.294 |
